# Neural dynamics of proactive and reactive cognitive control in medial and lateral prefrontal cortex

**DOI:** 10.1101/2025.02.12.637987

**Authors:** Anas U. Khan, Colin Hoy, Kristopher L. Anderson, Vitoria Piai, David King- Stephens, Kenneth D. Laxer, Peter Weber, Jack J. Lin, Robert T. Knight, J. Nicole Bentley

**Affiliations:** Department of Neurosurgery, University of Alabama at Birmingham, Birmingham, AL, USA; Department of Neurology, University of California, San Francisco, San Francisco, CA, USA; Helen Wills Neuroscience Institute, University of California, Berkeley, Berkeley, CA, USA; Radboud University, Donders Institute for Brain, Cognition, and Behaviour, Nijmegen, Netherlands; Department of Neurology and Neurosurgery, California Pacific Medical Center, San Francisco, CA, USA; Department of Neurology, University of California, Irvine, Irvine, CA, USA; Department of Neurology, University of California, Davis, Davis, CA, USA; Center for Mind and Brain, University of California, Davis, Davis, CA, USA; Departments of Psychology and Neuroscience, University of California, Berkeley, Berkeley, CA, USA

## Abstract

Goal-directed behavior requires adjusting cognitive control to both react to and prepare for conflict. Previous work indicates theta oscillations and population activity in dorsomedial prefrontal cortex (dmPFC) and dorsolateral prefrontal cortex (dlPFC) are critical for reactive control. However, the neural mechanisms supporting proactive control are less clear. Here, we investigated the neural basis of behavioral adaptations when control is prepared in anticipation of conflict using intracranial EEG (iEEG) in dmPFC and dlPFC during a Stroop task where conflict frequency was manipulated across blocks. We observed canonical conflict-driven increases in dmPFC theta and in dmPFC and dlPFC local population activity, as indexed by high frequency activity (HFA). Conflict also suppressed theta power in both regions after the response, accentuated a pre-response beta desynchronization selectively in dlPFC, and increased a post-response beta rebound in both regions. Importantly, we identified a pre-trial marker of proactive control where dmPFC theta power increased before trials when conflict was expected, and theta, beta, and HFA conflict signals in both regions were enhanced when conflict was rare and diminished when conflict was common. These findings reveal shared HFA but dissociable oscillatory dynamics in dmPFC and dlPFC during reactive conflict processing, highlight pre-trial dmPFC theta as a potential substrate for proactive control, and refine the roles of dmPFC and dlPFC in control adaptations.

## Introduction

Goal-directed behavior requires exerting cognitive control to resolve conflict between competing options, and control resources are strategically adjusted based on recent experience and future conflict expectations^1^. Extensive research in humans and non-human primates supports the proposal that dorsomedial prefrontal cortex (dmPFC) detects the need for control and recruits dorsolateral prefrontal cortex (dlPFC) to implement cognitive control^2–10^, with theta oscillations and local population activity enhanced in these key regions during conflict processing^6–8,10,11^. However, the mechanisms facilitating cognitive control adjustments in preparation for conflict remain unclear.

The dual mechanisms framework of cognitive control posits separate but complementary processes of proactive and reactive control^12^. Proactive control involves preparing cognitive resources for top-down control of behavior, whereas reactive control is triggered in response to control-demanding stimuli, such as conflict between automatic word reading and instructed color naming in the Stroop task^13^. Reactive control signals in dmPFC and dlPFC are well-studied during within-trial conflict processing, where local population activity in the dlPFC and dmPFC increases, as measured using fMRI BOLD signal^14,15^, single unit firing^5,10,16^, and intracranial EEG (iEEG) high frequency activity (HFA)^7,8^, which is a proxy of local multi-unit activity^17–19^. Conflict also increases theta oscillations in dmPFC, but not dlPFC^6,8^, which is proposed to reflect dmPFC’s role in detecting conflict and recruiting control resources^1,2,20–22^. Recent work also shows theta oscillations coordinate neural firing in human dmPFC and dlPFC during conflict, and this was also true for beta oscillations, which are proposed to gate information processing by increasing or decreasing to allow reinforcement or modification of neural states, respectively^10^.

Moreover, theta, beta, and population activity signals are linked to adaptations on trials after conflict, including response slowing (i.e., the Gratton effect^5,8,9,23,24^). However, these between-trial adaptations reflect a mixture of reactive control from the previous trial and proactive control preparations^25–27^. In sum, theta and population activity in dmPFC and dlPFC are critical control signals elicited by conflict that are implicated in short-timescale adaptations, but few studies have examined their role in proactive control.

Behavioral studies have isolated proactive control by increasing the frequency of difficult trials, which report performance improvements when conflict is more common (e.g., reduced conflict effects on reaction times (RTs))^24,28–31^. The dual mechanisms framework hypothesizes that stronger conflict expectations drive anticipatory allocation of cognitive resources^12,25^, and several scalp EEG studies have shown control-demanding trials elicit lower mid-frontal theta power when these trials are more frequent, consistent with reduced need for reactive, within-trial control signals^24,30,31^. One recent study showed pre-trial dmPFC and dlPFC single unit firing predicted conflict-related RTs, suggesting preparatory activity can influence conflict processing^32^. However, no study to date has elucidated a candidate mechanism for proactive control adaptations based on task demands, leaving the roles of dmPFC and dlPFC undefined.

Here, we address these issues by utilizing the high spatio-temporal resolution of iEEG recordings from human dmPFC and dlPFC during a Stroop task that manipulated conflict frequency across blocks. We hypothesized that stronger conflict expectations would reduce conflict effects on RTs and reactive neural activity in the dmPFC and dlPFC, and we also predicted that theta oscillations in dmPFC would track these proactive control adaptations. We first re-examined dmPFC and dlPFC activity during within-trial conflict processing, confirming canonical conflict-driven increases in dmPFC theta and HFA in both dmPFC and dlPFC, as well as providing additional insights into a shared post-response theta suppression and dissociable effects of conflict on beta power across regions. Manipulating conflict expectations uncovered an increase in pre-trial dmPFC theta power when conflict was common, revealing a potential neural substrate for proactive cognitive control. These results refine and expand the role of theta, beta, and local population activity in reactive and proactive control and elucidate the dissociable roles of dmPFC and dlPFC during goal-oriented behavior.

## Results

### Behavioral signatures of proactive and reactive control

iEEG activity in dmPFC and dlPFC was recorded while 23 people undergoing neurosurgical treatment for epilepsy performed a color-word Stroop task. Conflict expectations were manipulated by varying the percentage of congruent (%Cong) versus incongruent trials across blocks, but all blocks had a constant proportion of neutral trials (“XXXX”), allowing assessment of proactive control on post-neutral trials while controlling for previous trial type and frequency. We used linear mixed effects models (LMMs) to predict RTs based on task variables, revealing a main effect of current trial type. RTs were slower for Conflict (i.e., incongruent) trials, while congruent and neutral trials were not different and were thus combined into a NoConflict condition for subsequent analyses (see Methods) (n=23 subjects, 5888 trials, LMM, t_5860_=34.1, p=2×10^-16^; Figure 1D). We also found several between-trial adaptation effects to previous conflict, including post-conflict slowing (see Supplemental Information), but our primary focus was on proactive adaptations to %Cong conditions. We observed that NoConflict RTs were faster when conflict was rare (main effect of %Cong, t_5860_=-3.27, p=0.001), and that the effect of conflict on RTs was larger when conflict was more surprising (CurrentConflict:%Cong interaction, t_5860_=5.06, p=4.36×10^-07^; Figure 1E). In summary, the slowing effect of CurrentConflict was reduced in low %Cong blocks, indicating proactive control reduces the classic reactive Stroop effect when conflict is expected.

**Figure 1.**
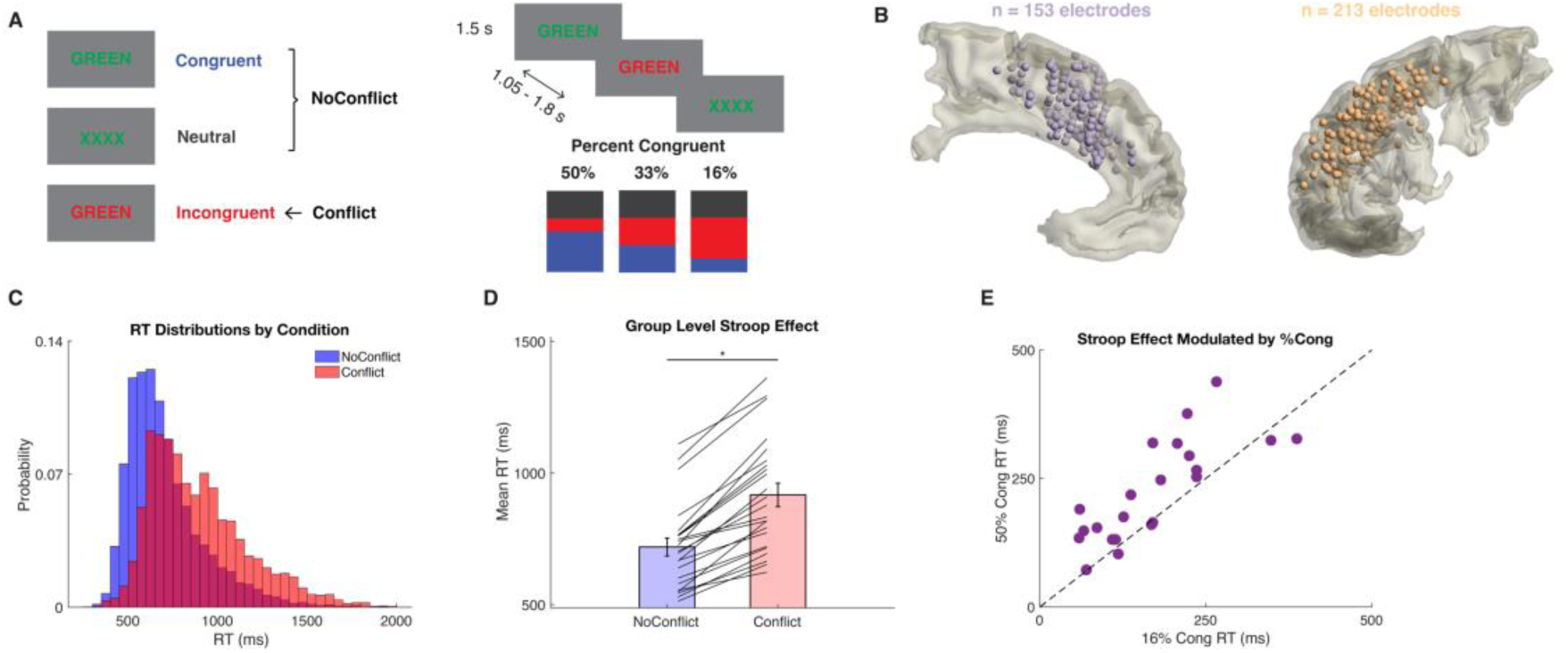
Task, behavior, and electrode locations. *A*, Trial types, %Cong block types, and trial design. *B*, Group-level reconstructions of recording sites for dmPFC and dlPFC. All sites mirrored to left hemisphere for visualization. *C*, RT distributions for Conflict and NoConflict trials. *D,* Stroop effect: RTs are longer on Conflict than NoConflict trials. *E*, Stroop effect (mean RT difference for Conflict-NoConflict) is larger when conflict is rare (16%Cong) than when conflict is common (50%Cong), indicating proactive control adaptation based on conflict expectations.

### Theta, beta, and HFA dynamics in dmPFC and dlPFC during within-trial conflict processing

To first establish the neural dynamics underlying reactive conflict processing within a trial, we used time-resolved LMMs to predict single-trial neural power in theta, beta, and HFA bands (Figure 2). In line with previous conflict studies ^6,8,9,24,33–35^, theta power was higher during response preparation in Conflict than NoConflict trials in dmPFC (LMM, all p<=0.022) but not dlPFC (Figure 2C). Similarly, dmPFC and dlPFC both showed HFA increases just before and during the response for Conflict relative to NoConflict trials (LMM, dmPFC: all p=0, dlPFC: all p<=0.006; Figure 2B). Both regions exhibited pre-response beta suppression, but only dlPFC showed a greater beta decrease on Conflict than NoConflict trials (LMM, all p<=0.008; Figure 2D), reflecting additional processing of conflict. Finally, dlPFC theta power showed a peri/post-response suppression in Conflict relative to NoConflict trials (LMM, p<=0.001; Figure 2C), which also was observed for dmPFC theta after the response (LMM, p<=0.004; Figure 2C). In summary, we observed a cascade of shared HFA but dissociable theta and beta dynamics in dmPFC and dlPFC, where conflict initially increased dmPFC theta and dlPFC HFA while suppressing dlPFC beta, then dmPFC HFA increased before a late suppression of theta in dlPFC then dmPFC.

**Figure 2.**
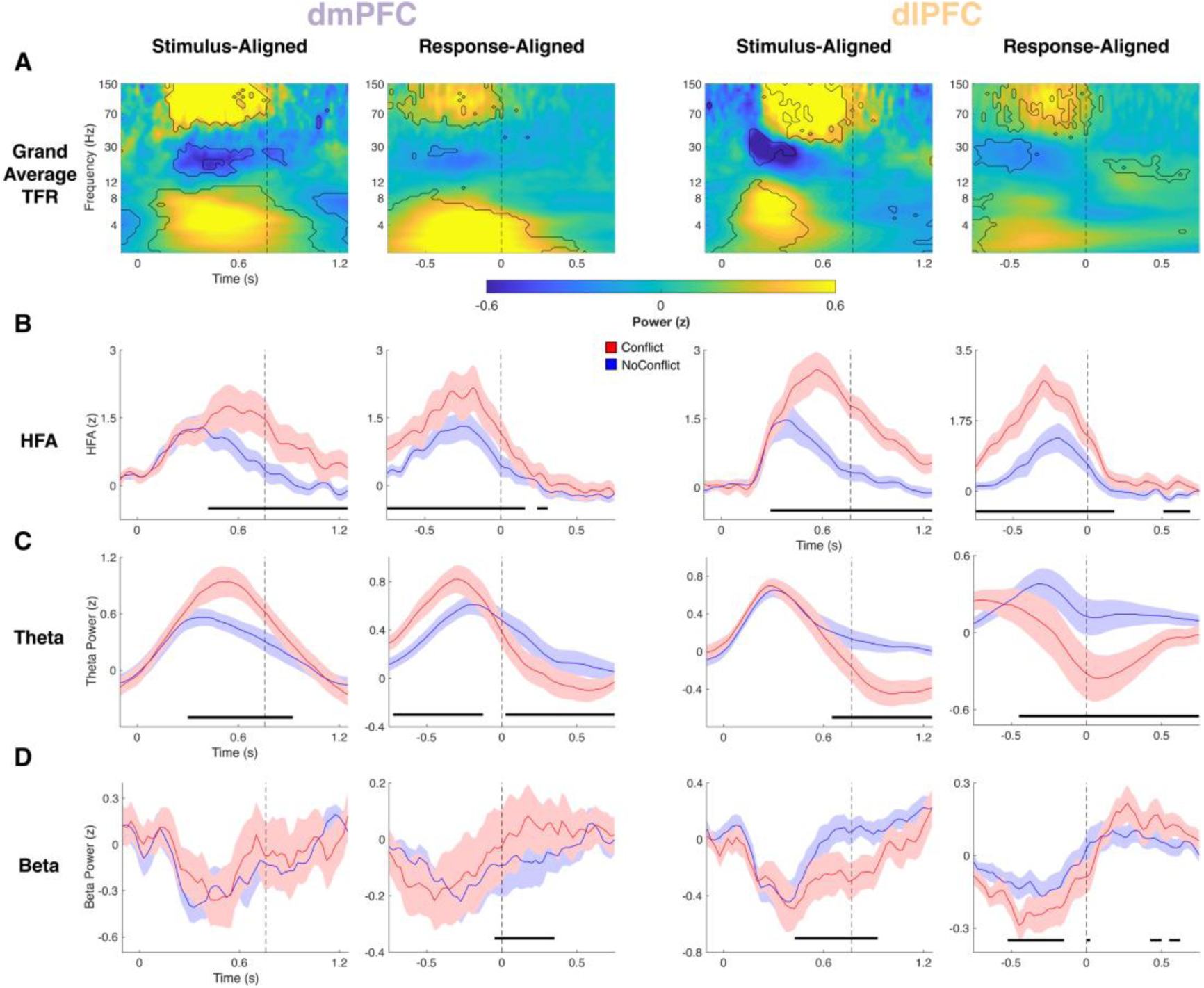
Conflict drives similar HFA and diverging theta and beta dynamics in dmPFC and dlPFC. *A*, Grand average time-frequency representations for dmPFC (Left pair) and dlPFC (Right pair) task encoding electrodes for both stimulus aligned (Left column of each region’s panel) and response-aligned (Right column of each region’s panel) data. Contour lines indicate power values significantly different from baseline. *B*, Time courses for HFA on Conflict (Red) and NoConflict (Blue) trials. *C*, Analogous time courses for theta power. *D*, Analogous time courses for beta power. Horizontal black lines indicate significance for CurrentConflict. Vertical dashed lines indicate group mean RT in stimulus-aligned data and RT in response aligned data.

### Conflict expectations modulate within-trial conflict processing and increase pre-trial theta power in dmPFC

To determine the neural basis of proactive control, we investigated how within-trial conflict responses were modulated by conflict anticipation. We found that theta and HFA responses in both regions were accentuated when conflict was rare and attenuated when conflict was expected (LMM, CurrentConflict:%Cong interaction, dmPFC: all p<=0.02, dlPFC: all p<=0.023; Figure 3A-D). However, the nature of these modulations varied across regions and frequency bands. In the theta band, we found an early dissociation where stronger expectations of conflict diminished responses in dmPFC but not dlPFC (CurrentConflict:%Cong interaction, LMM, all p<=0.011; Figure 3C-D). In contrast, both regions showed a larger effect of conflict on HFA when conflict was less expected. In the later post-response window, both regions showed a greater suppression of theta in blocks when conflict was rare (CurrentConflict:%Cong interaction, dmPFC: all p<=0.012, dlPFC: all p<=0.016; Figure 3C-D), suggesting theta during response monitoring is downregulated throughout the network when conflict resolution was more demanding. Pre-response beta power in dlPFC (but not dmPFC) was suppressed more for conflict trials when it was more surprising (i.e., less proactive control available) (CurrentConflict:%Cong interaction, all p<=0.022; Figure 3E-F).

**Figure 3.**
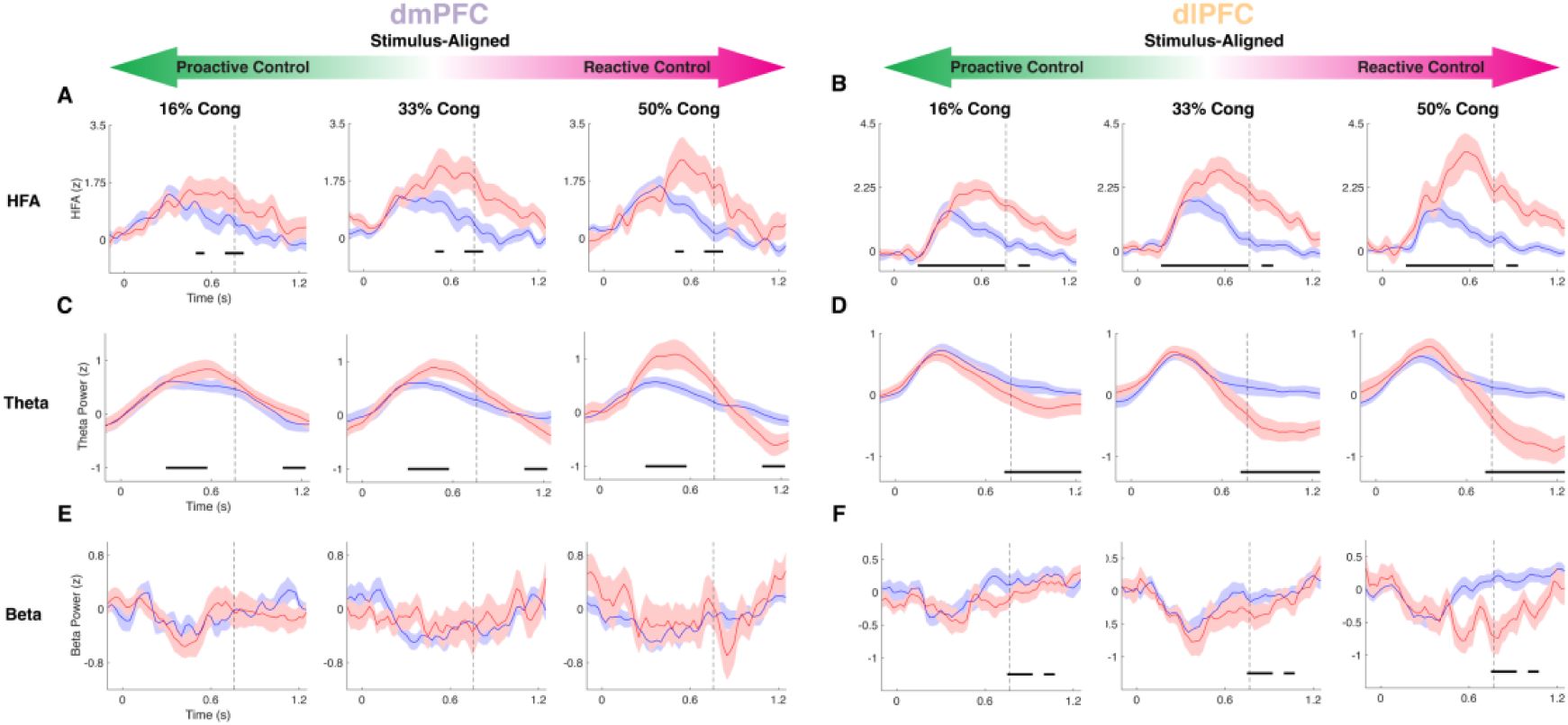
Bi-directional modulation of neural conflict signals by proactive control. *A*, dmPFC HFA in Conflict (red) and NoConflict (blue) trials across 3 block types, from highest to lowest proactive control (left to right: 16%, 33%, 50% congruent). *B,* Same as in *A* but for dlPFC. *C*, *D* Same as *A*, *B* but for theta power. *E*, *F* Same as above but for beta power. Horizontal black lines indicate significance for the CurrentConflict:%Cong interaction. Vertical dashed lines indicate group mean RT.

Finally, we tested our hypothesis that dmPFC theta facilitates these proactive control effects by examining %Cong block effects on neural activity in the preparatory period from-1s to stimulus onset. Notably, we avoided confounds of previous trial conflict and frequency by constraining this analysis to trials preceded only by neutral trials (see Methods). This revealed elevated dmPFC theta power approximately 0.5 s before stimulus onset in blocks when conflict was more frequent (LMM, p<=0.023, permutation test; Figure 4). In summary, higher proactive control reduced within-trial conflict dynamics and increased pre-trial dmPFC theta.

**Figure 4.**
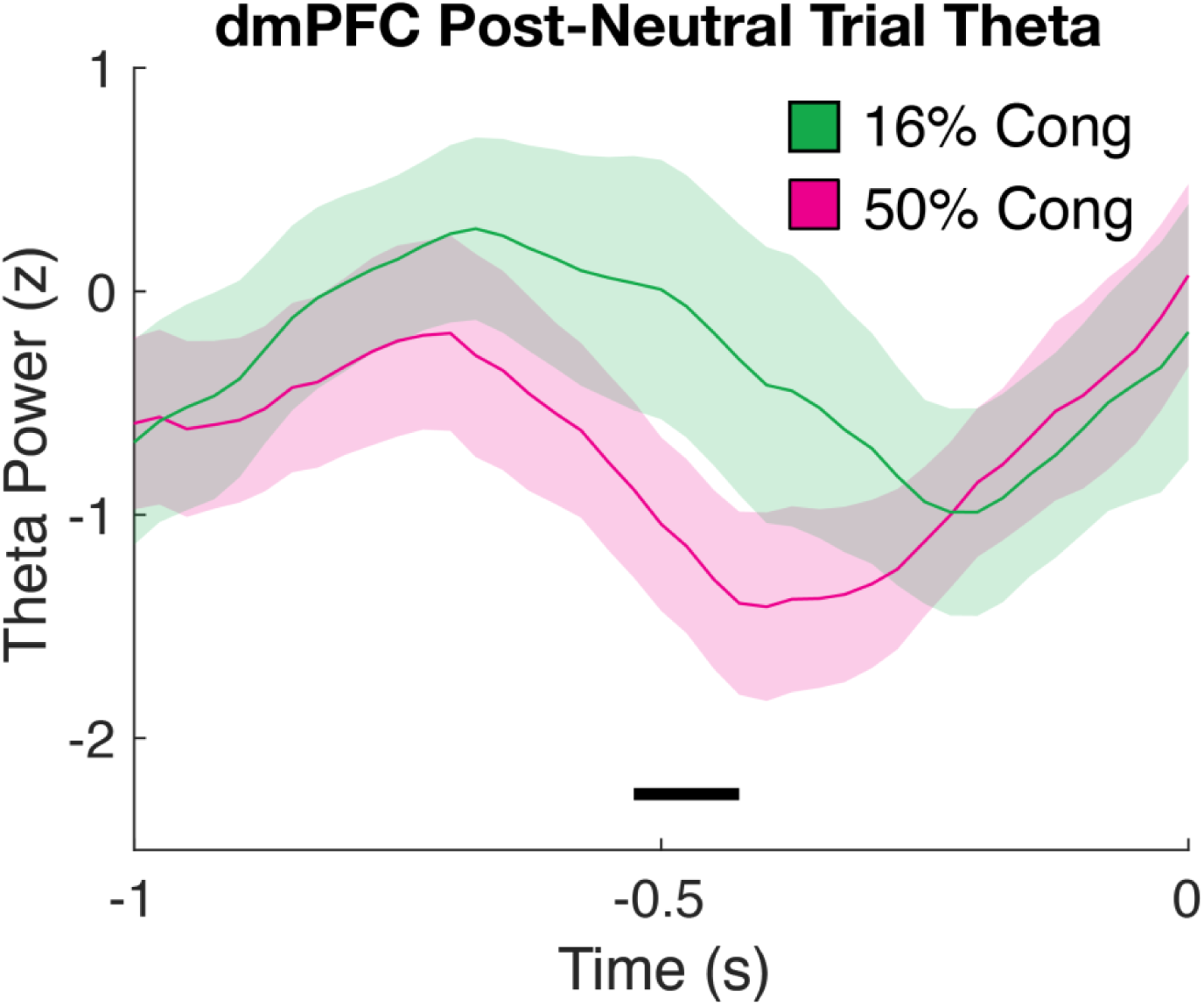
Proactive control increases pre-trial dmPFC theta power. Time courses for stimulus-aligned, post-neutral trial theta power in dmPFC for the 16% Congruent block (Green) and 50% Congruent block (Purple). Black line indicates significance for the %Cong effect.

## Discussion

We used iEEG in human dmPFC and dlPFC to investigate the neural dynamics of proactive and reactive cognitive control by manipulating conflict expectations in a Stroop task. Single-trial modeling of neural power revealed a sequence of within-trial conflict processing dynamics that differentiated dmPFC and dlPFC. In line with previous findings^6–9,24,33–35^, conflict triggered an increase in dmPFC theta and HFA increases in both regions. Building on these classic effects, we report novel effects of conflict, which increased a pre-response beta desynchronization in dlPFC and a post-response beta rebound in both regions, as well as suppressing theta in both regions after responses to conflict. Furthermore, these neural signals were accentuated when conflict was rare and diminished when conflict was common and thus expected, demonstrating how these within-trial dynamics are modulated by the balance between proactive and reactive control. Importantly, we also found that pre-trial theta power in the dmPFC was enhanced during blocks with strong expectation of conflict, revealing a novel signature of proactive control. Collectively, these results characterize how the cascade of neural signals in dmPFC and dlPFC during conflict processing are modulated by proactive and reactive control.

Our findings on the within-trial sequence of dmPFC and dlPFC neural signals underlying conflict processing corroborate and extend prior findings implicating these regions in cognitive control. Increases in HFA in both dmPFC and dlPFC during conflict suggest local neuronal populations within these regions are recruited to resolve competition between responses, aligning with earlier reports of dmPFC and dlPFC involvement in conflict detection and resolution^2,6–8,10,14^. However, the oscillatory dynamics in these regions diverged, providing insight into how dmPFC and dlPFC differentially contribute to conflict processing across different timescales.

Consistent with classical findings^2,5,14,36^, dmPFC showed robust increases in theta power during the response preparation epoch of Conflict compared to NoConflict trials, highlighting its role in conflict monitoring and signaling the need for control. In contrast, conflict triggered a strong post-response suppression of dlPFC theta power below baseline. A similar but weaker and later theta suppression after conflict was observed in dmPFC. This novel finding suggests that theta oscillations may be involved in resetting or down-regulating control circuits after a demanding event, which may help minimize control costs via efficient resource allocation. Overall, these findings suggest a temporally structured cascade of conflict processing across dmPFC and dlPFC. dmPFC theta and dlPFC HFA both rise with conflict, reflecting the concurrent detection of conflict (dmPFC) and engagement of control processes (dlPFC), with the subsequent increase in dmPFC HFA potentially reflecting ongoing conflict and performance monitoring. Taken together, these dynamics exemplify how dmPFC and dlPFC handle distinct roles in the cognitive control network.

We also observed dissociable effects of conflict on beta power across dmPFC and dlPFC. The pre-response decrease in beta power was stronger for Conflict than NoConflict trials in dlPFC but not dmPFC, whereas the post-response beta rebound increased on Conflict trials in both regions. Substantial research has linked dlPFC to maintenance and updating of working memory supporting task demands^14,37–39^, where the beta rhythm is proposed as a gating mechanism in these circuits with decreases in beta power opening a window for updating^39–41^.

Accordingly, the greater decrease in dlPFC beta on conflict trials may reflect additional processing demands required to exert control during conflict resolution^6,20^. In contrast, previous studies implicate post-response beta increases in regulating the integration of surprising feedback into an internal model of motor commands, where stronger beta increases indicate confidence in maintaining the current motor plans^39,42–45^. This suggests that the increased beta rebound in both regions after correct conflict responses may reflect reinforcement of the task rules. This effect was earlier in dmPFC than dlPFC, which aligns with proposals that dmPFC monitors control demands and relays this information to dlPFC^20^.

After establishing these theta, beta, and HFA dynamics during within-trial conflict processing, we addressed our main objective by examining how these dmPFC and dlPFC signals were modulated by proactive control as conflict expectations build up over time. Behaviorally, participants had a reduced Stroop effect when conflict was more expected, and as hypothesized, these different levels of proactive control revealed a bidirectional influence on neural conflict processing. Canonical increases in dmPFC theta and HFA in dmPFC and dlPFC on Conflict trials were exacerbated for high %Cong blocks but nearly absent in blocks when half of trials had conflict. Similarly, the novel suppression of pre-response dlPFC beta and post-response dmPFC and dlPFC theta on Conflict trials was larger after surprising conflict and showed a smaller effect after common conflict. Accentuated beta decreases in dlPFC during surprising conflict trials support interactions between reactive control and working memory, possibly reflecting larger working memory updates for flexible reconfiguration when conflict is rare. It remains unclear whether this effect reflects reactive, within-trial adjustments to the current conflict or facilitates updating conflict expectations on future trials. However, our findings underscore a key role for dlPFC beta in enabling goal-oriented behavior. Overall, this striking demonstration that theta, beta, and HFA conflict signals are exaggerated or attenuated when proactive control is minimal or maximal, respectively, refines the interpretation of these dynamics by linking them more closely to reactive control mechanisms.

Critically, we found that strong conflict expectations increased dmPFC theta power before trial onset, suggesting dmPFC theta may facilitate proactive control by preparing cognitive resources based on expected demands. Notably, this analysis was limited to post-neutral trials, and our task design held the rate of neutral trials constant across blocks, meaning this effect cannot be explained by reactive control adjustments to prior trial frequency or difficulty. Relatedly, post-conflict slowing predicted reduced early dmPFC theta (see Supplementary Information), implicating dmPFC theta in both between-trial and block-level control adaptations.

In sum, our results shed new light on how dmPFC and dlPFC conflict processing is modulated by proactive cognitive control adjustments. In addition to canonical theta and HFA increases to conflict, we show that conflict suppresses theta after the response in both dmPFC and dlPFC, potentially reflecting downregulation in control networks after intensive processing. We also report novel pre-response beta desynchronization in dlPFC and post-response beta rebound in dmPFC that align with the roles for these regions in exerting and monitoring control, respectively. Moreover, by varying task demands over long timescales, we show that stronger conflict expectations boost dmPFC theta prior to trial onset and reduce within-trial processing. Overall, these findings identify a putative mechanism by which proactive control improves performance by preparing cognitive resources to reduce reliance on reactive control.

## Methods

### Participants

Data was collected from 31 surgical epilepsy patients (mean ± SD = 35.7 ± 12.4 years old; 8 women). However, patients were excluded for technical problems during recordings (n = 4), poor data quality (n = 2), or lack of dmPFC or dlPFC coverage (n = 2). The 23 patients available for analyses were implanted with stereo-electroencephalography (SEEG; n = 21) probes and/or subdural electrocorticography (ECoG; n = 3) grids/strips. Patients were studied at the University of California, Berkeley, University of California, Irvine, and California Pacific Medical Center. All patients had normal IQ and spoke English as a primary language except two, which were both fluent in English with IQ above 85.

### Behavioral Task

Patients performed a color-word Stroop task written in PsychoPy^46^ (v1.82.01) with congruent, incongruent, and neutral stimuli consisting of the words “BLUE”, “RED”, “GREEN”, and “XXXX” (neutral) displayed in blue, red, or green ink on a gray computer screen (Fig. 1a). Color-word stimuli were presented for 1.5 s followed by inter-trial intervals randomly sampled from a uniform distribution between 1.05 and 1.8 s, except for one patient who had inter-trial intervals ranging from 2 to 2.3 s. Patients were asked to name the color of the ink, and verbal RTs were obtained from the microphone signal by manually marking the first deflection in the audio waveform above baseline noise for vocalizations naming the color. The task was organized into 9 blocks of 36 trials, except one patient who had 12 blocks of 24 trials. One patient performed the task twice. The proportion of congruent trials was manipulated across blocks such that they were either mostly congruent (50% congruent, 33.3% neutral, and 16.7% incongruent), equal proportions (33.3% congruent, 33.3% neutral, and 33.3% incongruent), or mostly incongruent (16.7% congruent, 33.3% neutral, and 50% incongruent; Figure 1A). Note that the consistent frequency of neutral trials across block types provides an unbiased diagnostic condition of block-level changes in conflict expectations. Block types were randomized and occurred in equal numbers.

### Intracranial EEG Recordings

Data were recorded at either the University of California (UC) Irvine Medical Center (n = 28), USA or California Pacific Medical Center (n = 3), USA. Patients at Irvine were implanted with SEEG electrodes with 5 mm spacing and/or ECoG grids with 1 cm spacing, and patients at CPMC were implanted with ECoG strips with 1 cm spacing. At both sites, electrophysiology and analog photodiode event channels were recorded using a 256-channel Nihon Kohden Neurofax EEG-1200 recording system and sampled at 500 (n = 2), 1000 (n = 2), or 5000 Hz (n = 19). For three datasets from UC Irvine, a separate Neuralynx ATLAS recording system was used to record the analog photodiode channels (n = 2 at 4000 Hz and n=1 at 8000 Hz) and a subset of iEEG channels (n = 1 at 4000 Hz and n = 2 at 8000 Hz). Photodiode events were then aligned to the iEEG data acquired in parallel via the Nihon Kohden clinical amplifier based on cross-correlation of shared iEEG channels.

### Electrode Localization

Pre-operative T1 MRI and post-implantation CT scans were collected as part of standard clinical care, and recording sites were reconstructed in patient space by aligning scans via rigid-body co-registration as described in Stolk et al.^47^. Anatomical locations of electrodes were determined by manual inspection in native patient space under supervision of a neurologist.

Electrode positions were then warped to a standard MNI 152 template brain using volume-based registration in SPM 12 as implemented in Fieldtrip^47^. Group-level electrode positions are plotted in MNI coordinates relative to the cortical surface of the fsaverage brain template from FreeSurfer^48^, with right hemisphere electrodes mirrored to visualize all electrodes on the left hemisphere.

### Data Preprocessing

Data cleaning, preprocessing, and analyses were conducted using the Fieldtrip toolbox^49^ and custom MATLAB code. Raw iEEG traces were manually inspected by a neurologist for epileptiform discharges along and artifacts (e.g., machine noise, signal drift, amplifier saturation, etc.). Data in regions or epochs with epileptiform or artifactual activity were excluded from further analyses. Preprocessing involved applying a 0.5-250 Hz anti-aliasing Butterworth filter and notch filtering for line noise at 60 Hz and at the next 4 harmonics (120, 180, 240, and 300 Hz) using a 2 Hz bandwidth Butterworth filter, resampling to 500 Hz, and re-referencing using adjacent bipolar montages for SEEG and common average schemes for ECOG grids/strips.

Continuous data were then visually re-inspected for quality. Trials were rejected for task interruptions and behavioral outliers (RTs missing, <0.3 s, >2.0 s, or >3 standard deviations from the patient mean), including errors and partial errors, which were too infrequent to analyze.

Finally, trials were segmented from-0.25 to 2.5 seconds relative to stimulus onset and rejected for excessive variance in the preprocessed time series or the differentiated preprocessed time series. Exclusion criteria for trial variance were based on patient-specific thresholds of trial-level standard deviations ranging from 5 to 10 standard deviations. Between 0 and 13 trials per patient were rejected for excessive variance (mean ± S.D.: 4.0 ± 3.2 trials). In total, this process resulted in 128-605 trials per patient (mean ± S.D.: 282.5 ± 83.8) available for analyses.

### Time-Frequency Analysis

We used the FieldTrip toolbox^49^ and custom MATLAB (Mathworks) scripts to convolve trial data (S-locked:-0.5 to 1.25 s; R-locked:-0.75 to 0.75 s) with 51 logarithmically spaced complex Morlet wavelets (number of cycles = 6) ranging from 2 to 152 Hz. We included a 1.5 s buffer period on both sides of the signal, which was discarded afterwards. We obtained a continuous estimate of instantaneous power by squaring the magnitude of the continuous wavelet transform and log transformed the result in preparation for linear modeling. We then normalized the log-transformed power values by randomly sampling values from the baseline period (-0.5 to-0.2s relative to stimulus) of all trials 1000 times and used the mean and standard deviation of this surrogate distribution to z-score within each electrode and frequency separately. For the case of investigating the preparatory period, we normalized the power values to a bootstrapped distribution sampled from the entire task recording. HFA was treated as a separate aggregate band (70 - 150 Hz). It was created by bandpass filtering the time series in 8 10-Hz bins (70 - 80, 80 - 90, etc.) and extracting the amplitude of the Hilbert transform within each bin. The HFA was normalized in the same manner as described above, separately within each bin, then averaged together to produce one composite band. Theta (4 – 8 Hz) and beta (12 – 30 Hz) power was extracted by averaging single-trial data within the respective bands and downsampling to 40 Hz prior to statistical analysis to improve computational efficiency. HFA was downsampled to 100 Hz.

### Statistical Analysis

For all of our analyses, we used linear mixed-effects models (LMM). Neutral and Congruent trials were combined into “NoConflict” due to the lack of behavioral and neural differences between these conditions. We began with the omnibus model predicting RTs (Equation 1) and excluded terms that were not significant at α = 0.05 to reduce the model complexity.

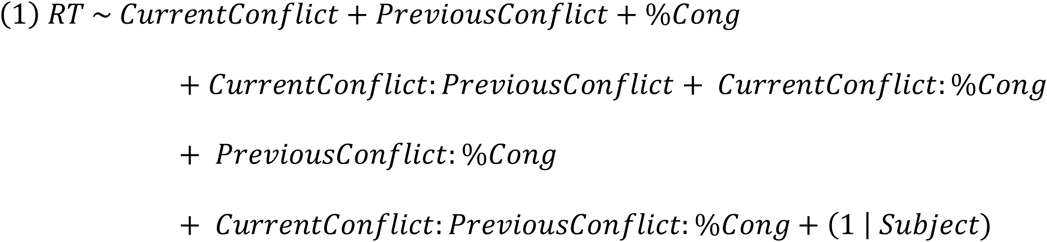

Here, CurrentConflict has 2 levels (Conflict and NoConflict) and represents the fixed effect of the current trial type, with NoConflict as the reference. PreviousConflict is the analogous factor for the trial type before the current trial. %Cong (percent congruent) is either 16%, 33%, or 50% and was centered at 33% in the model. Colons (“:”) indicate interactions and the “(1 | Subject)” term represents a random intercept for each subject. *P*-values were obtained using Wald *t* tests. The final model after the top-down model selection procedure is given below (Equation 2).

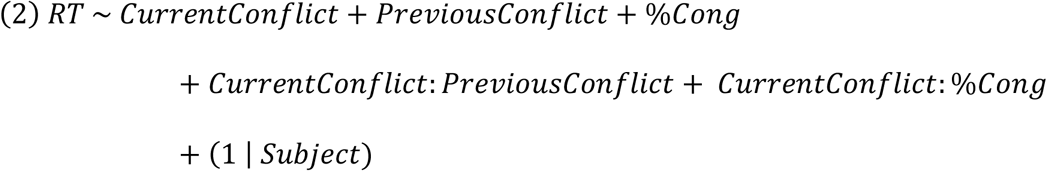

For neural data, we used a well-established information theoretic approach^47,50–54^ to select for task encoding electrodes for subsequent analyses. Task encoding was defined as an electrode in which trial type (con, neu, inc) explained a significant amount of variance in the HFA using a one-way analysis of variance (ANOVA). Electrodes that exhibited a significant main effect of trial type for more than 100 ms were kept for further analysis. This process yielded 93 task encoding electrodes for dmPFC and 121 for dlPFC. LMMs were performed at every time point. We corrected for multiple comparisons using permutation tests on the Wald *t* time series and applying threshold-free cluster enhancement (TFCE) with E = 0.66 and H = 2/3^55,56^. TFCE scores in the top 5% of the surrogate distributions were considered statistically significant. P values for permutation tests are reported as the largest p value among significant clusters for each effect.

## Funding Acknowledgements

This work was funded by NINDS R01NS021135, NIMH CONTE Center P50MH109429, NIMH F32MH132174, Welcome Discovery Award (226645/Z/22/Z), and NINDS 5K23NS117735.

## Supplemental Introduction

Cognitive control adjustments are commonly studied through between-trial adaptations. Foundational fMRI work demonstrates dmPFC activity on conflict trials predicts behavioral adjustments on the next trial^1,2^, the next trial’s adjustments correspond with greater dlPFC activation^1,2^, and this dlPFC activation is correlated with the previous trial’s dmPFC activation^1^. Recent human iEEG studies found that neurons in the anterior cingulate fire faster after conflict than after non-conflict trials^3^, and post-conflict dlPFC sustained HFA has been shown to correlate with longer reaction times (RTs)^4^. Although between-trial adaptations cannot disentangle the contributions of proactive and reactive control and are thus not the core focus of this study, we sought to connect our main findings to prior results by reporting supplemental behavioral and neural results on post-conflict adaptations.

## Supplemental Results

### Between-trial behavioral adaptations after conflict

We found a main effect of previous trial conflict, where RTs were slower after Conflict trials compared to after NoConflict trials (t_5860_=5.63, p=1.90 x 10^-08^). The interaction between current and previous conflict was also significant (t_5860_=-2.19, p=0.03), a phenomenon described previously as the Gratton effect ^5^. Post-hoc, pairwise comparisons revealed that the Stroop effect was largest when the previous trial was NoConflict (t_5860_=34.3, p=0, Tukey corrected), with a smaller but still significant effect when the previous trial was Conflict (t_5860_=20.3, p=2.15 x 10^-12^). The Gratton effect was driven by slower RTs on current NoConflict trials when the previous trial was Conflict (t_5860_=5.63, p=1.14 x 10^-07^, Figure S1), whereas Conflict trials were not affected by previous trial conflict (t_5860_=1.32, p=0.549).

### dmPFC theta tracks post-conflict slowing

We examined the neural basis of these within-trial reactive control dynamics by examining the neural basis of post-conflict slowing on NoConflict trials. While we did not find condition-specific differences in neural activity in either frequency band or either region based on previous Conflict versus NoConflict trials, we found a relationship between theta power and RT adjustments that was specific to dmPFC and the condition with behavioral slowing after conflict. Specifically, higher RTs (greater adjustments) predicted lower dmPFC theta power in the early stage of NoConflict trials following Conflict trials (LMM, all p<=0.021, Figure S2A).

Importantly, this effect of RTs on theta was absent on post-NoConflict trials without control adaptations (LMM, all p>=0.05, Figure S2C), and no RT adjustment effects were observed in dlPFC theta (Figure S2B, D). During the later, peri-response window, dmPFC theta power showed a general relationship with RTs on both post-Conflict and post-NoConflict trials, where trials with longer RTs exhibited greater theta power (LMM, all p<=0.01, Figure S2A, C), which is consistent with proposals linking dmPFC theta to reactive monitoring of control demands^6^.

### Response-locked oscillatory and population dynamics vary with conflict anticipation

To determine whether the theta, beta, and HFA conflict signals were best interpreted as stimulus-or response-driven, we repeated our main analysis investigating the interplay between proactive and reactive control by realigning our data to the response. We largely found consistent effects with stimulus-aligned data except for dmPFC theta and HFA, both of which were trending and visually apparent but did not reach statistical significance (Figure S3).

## Supplemental Discussion

On the trial-by-trial timescale, we observed classical post-conflict slowing on NoConflict trials^5^, which is typically interpreted as increased caution after difficult responses. Ultimately, we did not find condition-specific differences in neural power related to previous trial type.

Instead, we found that early trial dmPFC theta power was reduced on NoConflict trials with stronger post-conflict slowing, which was specific to this region and condition. This reduction in the early dmPFC theta response may reflect a reduced need to recruit control resources due to prior recruitment on previous Conflict trials. Regardless, the selectivity of this relationship to conditions showing trial-by-trial behavioral adjustments aligns with our finding that pre-trial dmPFC theta increases with proactive control to support the theory that dmPFC theta power plays a key role in managing control over multiple time scales^7^.

---

**Figure S1.**
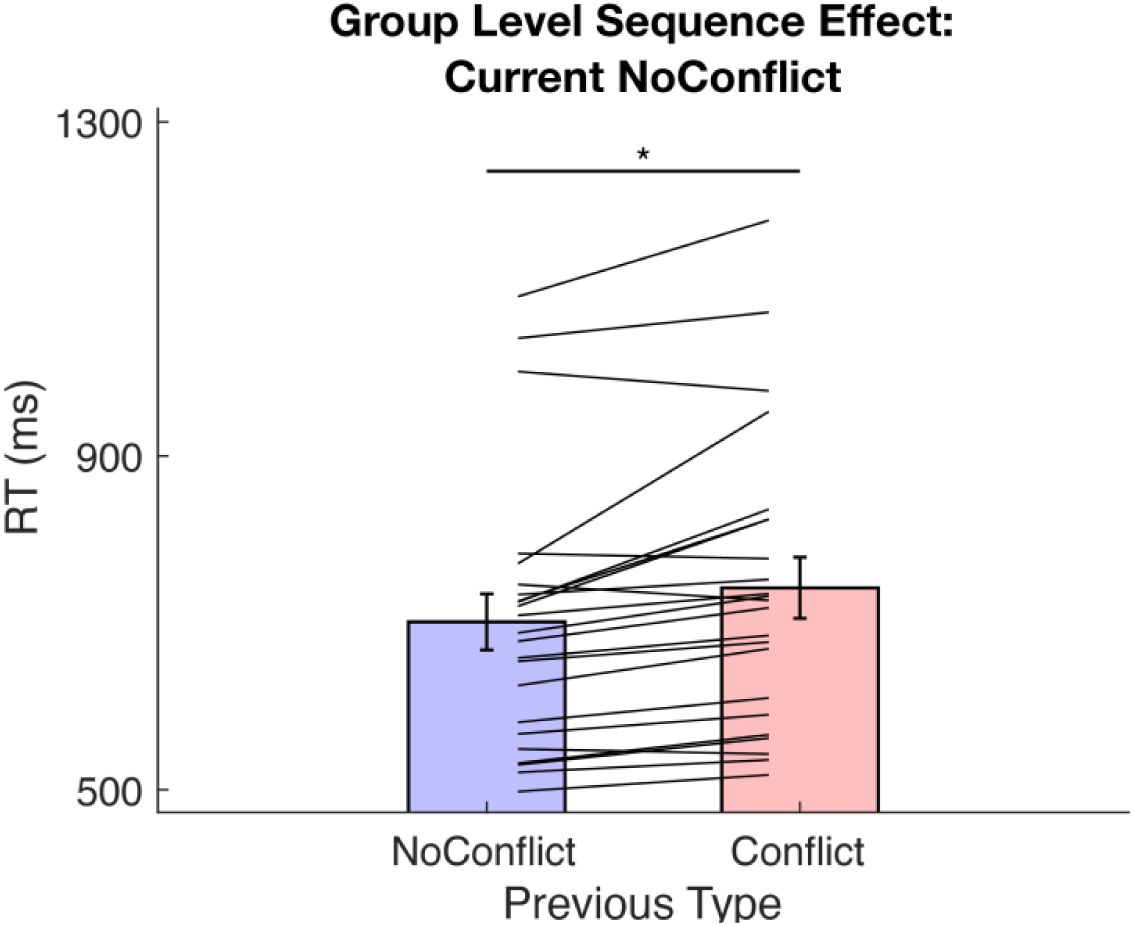
Conflict prolongs response times on subsequent NoConflict Trials. Bar graphs depicting mean RT across subjects with individual subject mean RTs shown as lines. RTs were taken only from current NoConflict trials and plotted as a function of previous conflict.

**Figure S2.**
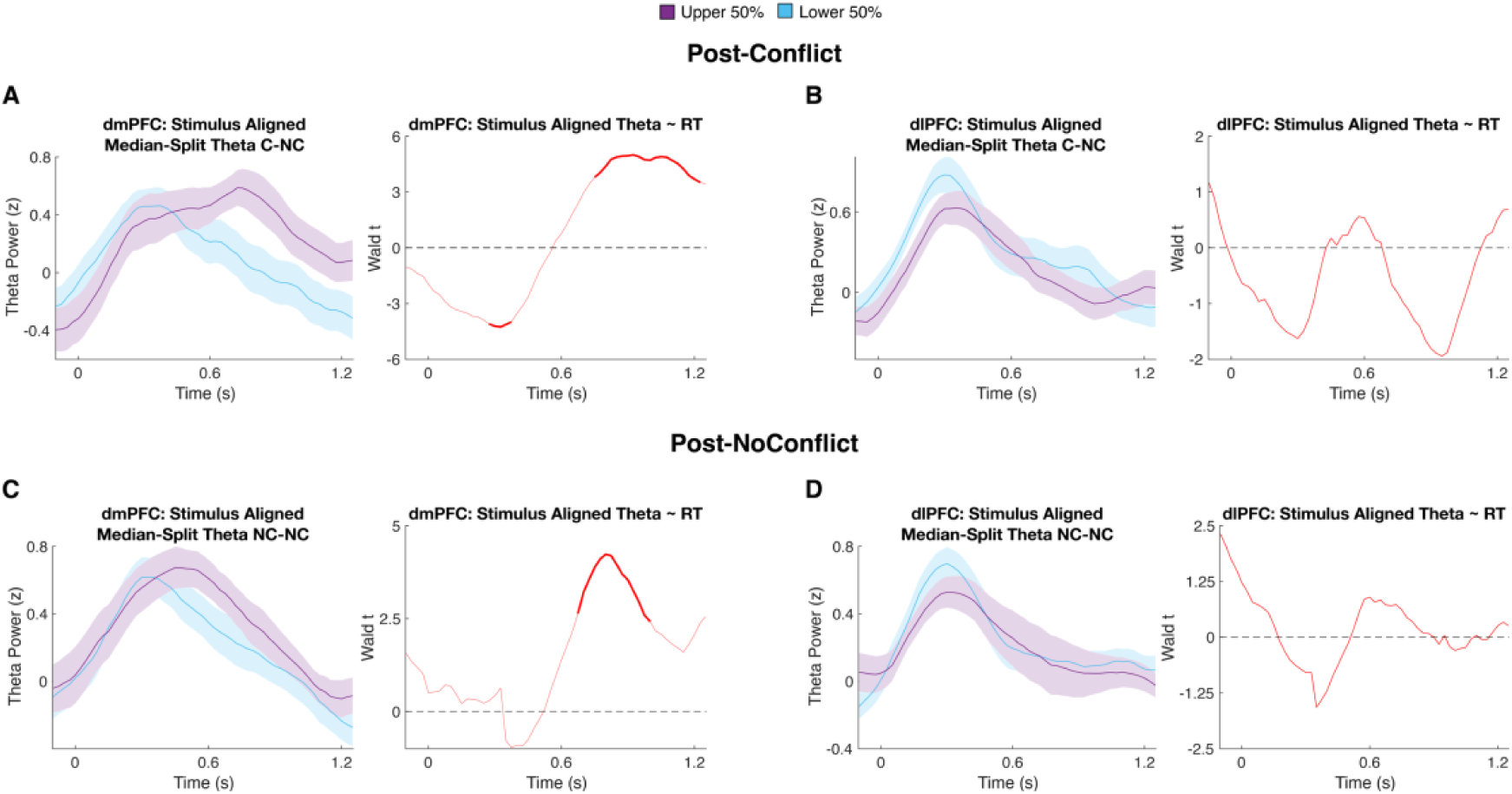
dmPFC theta power indexes between-trial control adjustments. *A*, (Left) Stimulus-locked dmPFC theta power averaged for slow (purple) and fast (blue) RTs based on a median split of RTs on NoConflict trials preceded by Conflict. Statistics are based on LMM regressions, median split for visualization purposes only. (Right) Wald t time course from the LMM with RT predicting theta power on Conflict->NoConflict trials. *B*, Same as in *A* but for dlPFC. *C*, Same as in *A* but for NoConflict->NoConflict trials. *D*, Same as in *C* but for dlPFC.

**Figure S3.**
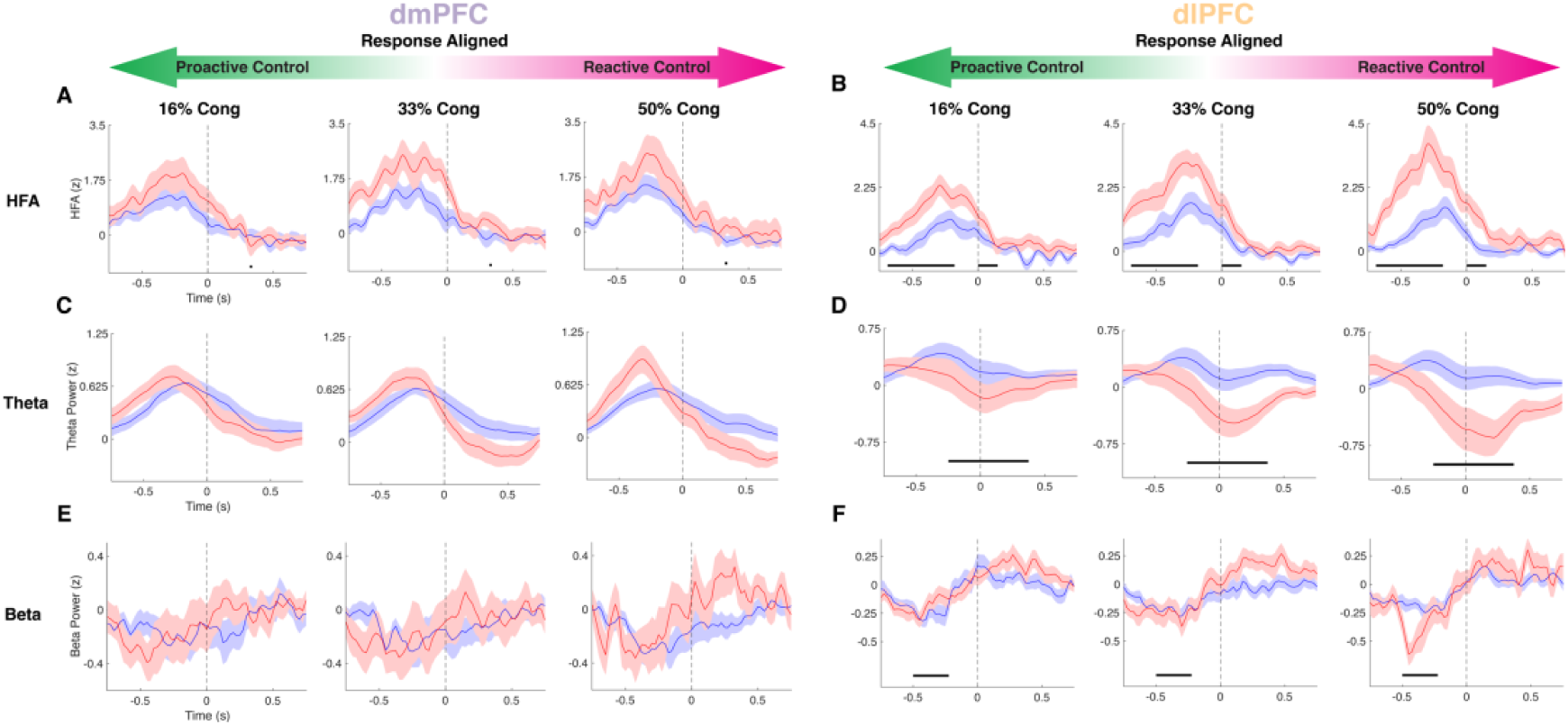
Bi-directional modulation of response-locked neural conflict signals by proactive control. *A*, dmPFC HFA in Conflict (red) and NoConflict (blue) trials across 3 block types, from highest to lowest proactive control (left to right: 16%, 33%, 50% congruent). *B,* Same as in *A* but for dlPFC. *C*, *D* Same as *A*, *B* but for theta power. *E*, *F* Same as above but for beta power. Horizontal black lines indicate significance for the CurrentConflict:%Cong interaction. Vertical dashed lines indicate the RT.

## References

1. Botvinick, M.M., Braver, T.S., Barch, D.M., Carter, C.S., and Cohen, J.D. (2001). Conflict monitoring and cognitive control. Psychol Rev 108, 624–652. 10.1037/0033-295x.108.3.624.

2. Kerns, J.G., Cohen, J.D., MacDonald, A.W., Cho, R.Y., Stenger, V.A., and Carter, C.S. (2004). Anterior Cingulate Conflict Monitoring and Adjustments in Control. Science 303, 1023–1026. 10.1126/science.1089910.

3. Kerns, J.G. (2006). Anterior cingulate and prefrontal cortex activity in an FMRI study of trial-to-trial adjustments on the Simon task. NeuroImage 33, 399–405. 10.1016/j.neuroimage.2006.06.012.

4. Nee, D.E., Wager, T.D., and Jonides, J. (2007). Interference resolution: insights from a meta-analysis of neuroimaging tasks. Cogn Affect Behav Neurosci 7, 1–17. 10.3758/cabn.7.1.1.

5. Sheth, S.A., Mian, M.K., Patel, S.R., Asaad, W.F., Williams, Z.M., Dougherty, D.D., Bush, G., and Eskandar, E.N. (2012). Human Dorsal Anterior Cingulate Cortex Neurons Mediate Ongoing Behavioral Adaptation. Nature 488, 218–221. 10.1038/nature11239.

6. Oehrn, C.R., Hanslmayr, S., Fell, J., Deuker, L., Kremers, N.A., Do Lam, A.T., Elger, C.E., and Axmacher, N. (2014). Neural communication patterns underlying conflict detection, resolution, and adaptation. J Neurosci 34, 10438–10452. 10.1523/JNEUROSCI.3099-13.2014.

7. Tang, H., Yu, H.-Y., Chou, C.-C., Crone, N.E., Madsen, J.R., Anderson, W.S., and Kreiman, G. (2016). Cascade of neural processing orchestrates cognitive control in human frontal cortex. Elife 5, e12352. 10.7554/eLife.12352.

8. Bartoli, E., Conner, C.R., Kadipasaoglu, C.M., Yellapantula, S., Rollo, M.J., Carter, C.S., and Tandon, N. (2018). Temporal Dynamics of Human Frontal and Cingulate Neural Activity During Conflict and Cognitive Control. Cereb Cortex 28, 3842–3856. 10.1093/cercor/bhx245.

9. Zavala, B., Jang, A., Trotta, M., Lungu, C.I., Brown, P., and Zaghloul, K.A. (2018). Cognitive control involves theta power within trials and beta power across trials in the prefrontal-subthalamic network. Brain 141, 3361– 3376. 10.1093/brain/awy266.

10. Smith, E.H., Horga, G., Yates, M.J., Mikell, C.B., Banks, G.P., Pathak, Y.J., Schevon, C.A., McKhann, G.M., Hayden, B.Y., Botvinick, M.M., et al. (2019). Widespread temporal coding of cognitive control in the human prefrontal cortex. Nat Neurosci 22, 1883–1891. 10.1038/s41593-019-0494-0.

11. Fu, Z., Sajad, A., Errington, S.P., Schall, J.D., and Rutishauser, U. (2023). Neurophysiological mechanisms of error monitoring in human and non-human primates. Nat Rev Neurosci 24, 153–172. 10.1038/s41583-022-00670-w.

12. Braver, T.S. (2012). The variable nature of cognitive control: A dual-mechanisms framework. Trends Cogn Sci 16, 106–113. 10.1016/j.tics.2011.12.010.

13. Stroop, J.R. (1935). Studies of interference in serial verbal reactions. Journal of Experimental Psychology 18, 643–662. 10.1037/h0054651.

14. MacDonald, A.W., Cohen, J.D., Stenger, V.A., and Carter, C.S. (2000). Dissociating the Role of the Dorsolateral Prefrontal and Anterior Cingulate Cortex in Cognitive Control. Science 288, 1835–1838. 10.1126/science.288.5472.1835.

15. 15. van Veen, V., Cohen, J.D., Botvinick, M.M., Stenger, V.A., and Carter, C.S. (2001). Anterior cingulate cortex, conflict monitoring, and levels of processing. Neuroimage 14, 1302–1308. 10.1006/nimg.2001.0923.

16. Fu, Z., Wu, D.-A.J., Ross, I., Chung, J.M., Mamelak, A.N., Adolphs, R., and Rutishauser, U. (2019). Single-neuron correlates of error monitoring and post-error adjustments in human medial frontal cortex. Neuron 101, 165–177.e5. 10.1016/j.neuron.2018.11.016.

17. Ray, S., and Maunsell, J.H.R. (2011). Different Origins of Gamma Rhythm and High-Gamma Activity in Macaque Visual Cortex. PLoS Biol 9, e1000610. 10.1371/journal.pbio.1000610.

18. Rich, E.L., and Wallis, J.D. (2017). Spatiotemporal dynamics of information encoding revealed in orbitofrontal high-gamma. Nat Commun 8, 1139. 10.1038/s41467-017-01253-5.

19. Leszczyński, M., Barczak, A., Kajikawa, Y., Ulbert, I., Falchier, A.Y., Tal, I., Haegens, S., Melloni, L., Knight, R.T., and Schroeder, C.E. (2020). Dissociation of broadband high-frequency activity and neuronal firing in the neocortex. Sci Adv 6, eabb0977. 10.1126/sciadv.abb0977.

20. Shenhav, A., Botvinick, M.M., and Cohen, J.D. (2013). The expected value of control: An integrative theory of anterior cingulate cortex function. Neuron 79, 217–240. 10.1016/j.neuron.2013.07.007.

21. Botvinick, M.M., Cohen, J.D., and Carter, C.S. (2004). Conflict monitoring and anterior cingulate cortex: an update. Trends in Cognitive Sciences 8, 539–546. 10.1016/j.tics.2004.10.003.

22. Botvinick, M., Nystrom, L.E., Fissell, K., Carter, C.S., and Cohen, J.D. (1999). Conflict monitoring versus selection-for-action in anterior cingulate cortex. Nature 402, 179–181. 10.1038/46035.

23. Gratton, G., Coles, M.G., and Donchin, E. (1992). Optimizing the use of information: strategic control of activation of responses. J Exp Psychol Gen 121, 480–506. 10.1037//0096-3445.121.4.480.

24. Chinn, L.K., Pauker, C.S., and Golob, E.J. (2018). Cognitive control and midline theta adjust across multiple timescales. Neuropsychologia 111, 216–228. 10.1016/j.neuropsychologia.2018.01.031.

25. Braem, S., Bugg, J.M., Schmidt, J.R., Crump, M.J.C., Weissman, D.H., Notebaert, W., and Egner, T. (2019). Measuring adaptive control in conflict tasks. Trends Cogn Sci 23, 769–783. 10.1016/j.tics.2019.07.002.

26. Egner, T. (2007). Congruency sequence effects and cognitive control. Cognitive, Affective, & Behavioral Neuroscience 7, 380–390. 10.3758/CABN.7.4.380.

27. Van Den Wildenberg, W.P.M., Wylie, S.A., Forstmann, B.U., Burle, B., Hasbroucq, T., and Ridderinkhof, K.R. (2010). To Head or to Heed? Beyond the Surface of Selective Action Inhibition: A Review. Front. Hum. Neurosci. 4. 10.3389/fnhum.2010.00222.

28. Risko, E.F., Blais, C., Stolz, J.A., and Besner, D. (2008). Nonstrategic contributions to putatively strategic effects in selective attention tasks. J Exp Psychol Hum Percept Perform 34, 1044–1052. 10.1037/0096-1523.34.4.1044.

29. Torres-Quesada, M., Lupiáñez, J., Milliken, B., and Funes, M.J. (2014). Gradual proportion congruent effects in the absence of sequential congruent effects. Acta Psychol (Amst) 149, 78–86. 10.1016/j.actpsy.2014.03.006.

30. Dippel, G., Chmielewski, W., Mückschel, M., and Beste, C. (2016). Response mode-dependent differences in neurofunctional networks during response inhibition: an EEG-beamforming study. Brain Struct Funct 221, 4091–4101. 10.1007/s00429-015-1148-y.

31. Dippel, G., Mückschel, M., Ziemssen, T., and Beste, C. (2017). Demands on response inhibition processes determine modulations of theta band activity in superior frontal areas and correlations with pupillometry – Implications for the norepinephrine system during inhibitory control. NeuroImage 157, 575–585. 10.1016/j.neuroimage.2017.06.037.

32. Herman, A.B., Smith, E.H., Schevon, C.A., Yates, M.J., McKhann, G.M., Botvinick, M., Hayden, B.Y., and Sheth, S.A. (2023). Pretrial predictors of conflict response efficacy in the human prefrontal cortex. iScience 26, 108047. 10.1016/j.isci.2023.108047.

33. Cohen, M.X., and Cavanagh, J.F. (2011). Single-Trial Regression Elucidates the Role of Prefrontal Theta Oscillations in Response Conflict. Front Psychol 2, 30. 10.3389/fpsyg.2011.00030.

34. Cohen, M.X., and Donner, T.H. (2013). Midfrontal conflict-related theta-band power reflects neural oscillations that predict behavior. J Neurophysiol 110, 2752–2763. 10.1152/jn.00479.2013.

35. Cavanagh, J.F., and Frank, M.J. (2014). Frontal theta as a mechanism for cognitive control. Trends Cogn Sci 18, 414–421. 10.1016/j.tics.2014.04.012.

36. Cavanagh, J.F., Zambrano-Vazquez, L., and Allen, J.J.B. (2012). Theta Lingua Franca: A Common Mid-Frontal Substrate for Action Monitoring Processes. Psychophysiology 49, 220–238. 10.1111/j.1469-8986.2011.01293.x.

37. Swann, N.C., Tandon, N., Pieters, T.A., and Aron, A.R. (2013). Intracranial Electroencephalography Reveals Different Temporal Profiles for Dorsal-and Ventro-lateral Prefrontal Cortex in Preparing to Stop Action. Cereb Cortex 23, 2479–2488. 10.1093/cercor/bhs245.

38. Buschman, T.J., Denovellis, E.L., Diogo, C., Bullock, D., and Miller, E.K. (2012). Synchronous Oscillatory Neural Ensembles for Rules in the Prefrontal Cortex. Neuron 76, 838–846. 10.1016/j.neuron.2012.09.029.

39. Khan, A.U., Irwin, Z., Mahavadi, A., Roller, A., Goodman, A.M., Guthrie, B.L., Visscher, K., Knight, R.T., Walker, H.C., and Bentley, J.N. (2024). Low-Frequency Oscillations in Mid-rostral Dorsolateral Prefrontal Cortex Support Response Inhibition. J Neurosci 44, e0122242024. 10.1523/JNEUROSCI.0122-24.2024.

40. Spitzer, B., and Haegens, S. (2017). Beyond the Status Quo: A Role for Beta Oscillations in Endogenous Content (Re)Activation. eNeuro 4, ENEURO.0170-17.2017. 10.1523/ENEURO.0170-17.2017.

41. Engel, A.K., and Fries, P. (2010). Beta-band oscillations--signalling the status quo? Curr Opin Neurobiol 20, 156–165. 10.1016/j.conb.2010.02.015.

42. Schmidt, R., Ruiz, M.H., Kilavik, B.E., Lundqvist, M., Starr, P.A., and Aron, A.R. (2019). Beta Oscillations in Working Memory, Executive Control of Movement and Thought, and Sensorimotor Function. J. Neurosci. 39, 8231–8238. 10.1523/JNEUROSCI.1163-19.2019.

43. 43. Koelewijn, T., van Schie, H.T., Bekkering, H., Oostenveld, R., and Jensen, O. (2008). Motor-cortical beta oscillations are modulated by correctness of observed action. NeuroImage 40, 767–775. 10.1016/j.neuroimage.2007.12.018.

44. Tan, H., Wade, C., and Brown, P. (2016). Post-Movement Beta Activity in Sensorimotor Cortex Indexes Confidence in the Estimations from Internal Models. J Neurosci 36, 1516–1528. 10.1523/JNEUROSCI.3204-15.2016.

45. Torrecillos, F., Alayrangues, J., Kilavik, B.E., and Malfait, N. (2015). Distinct Modulations in Sensorimotor Postmovement and Foreperiod β-Band Activities Related to Error Salience Processing and Sensorimotor Adaptation. J Neurosci 35, 12753–12765. 10.1523/JNEUROSCI.1090-15.2015.

46. Peirce, J.W. (2008). Generating Stimuli for Neuroscience Using PsychoPy. Front Neuroinform 2, 10. 10.3389/neuro.11.010.2008.

47. Stolk, A., Griffin, S., van der Meij, R., Dewar, C., Saez, I., Lin, J.J., Piantoni, G., Schoffelen, J.-M., Knight, R.T., and Oostenveld, R. (2018). Integrated analysis of anatomical and electrophysiological human intracranial data. Nat Protoc 13, 1699–1723. 10.1038/s41596-018-0009-6.

48. Dale, A.M., Fischl, B., and Sereno, M.I. (1999). Cortical Surface-Based Analysis: I. Segmentation and Surface Reconstruction. NeuroImage 9, 179–194. 10.1006/nimg.1998.0395.

49. Oostenveld, R., Fries, P., Maris, E., and Schoffelen, J.-M. (2011). FieldTrip: Open source software for advanced analysis of MEG, EEG, and invasive electrophysiological data. Comput Intell Neurosci 2011, 156869. 10.1155/2011/156869.

50. Siegel, M., Buschman, T.J., and Miller, E.K. (2015). Cortical Information Flow During Flexible Sensorimotor Decisions. Science 348, 1352–1355. 10.1126/science.aab0551.

51. Voytek, B., Kayser, A.S., Badre, D., Fegen, D., Chang, E.F., Crone, N.E., Parvizi, J., Knight, R.T., and D’Esposito, M. (2015). Oscillatory dynamics coordinating human frontal networks in support of goal maintenance. Nat Neurosci 18, 1318–1324. 10.1038/nn.4071.

52. Dürschmid, S., Edwards, E., Reichert, C., Dewar, C., Hinrichs, H., Heinze, H.-J., Kirsch, H.E., Dalal, S.S., Deouell, L.Y., and Knight, R.T. (2016). Hierarchy of prediction errors for auditory events in human temporal and frontal cortex. Proc Natl Acad Sci U S A 113, 6755–6760. 10.1073/pnas.1525030113.

53. Weber, J., Iwama, G., Solbakk, A.-K., Blenkmann, A.O., Larsson, P.G., Ivanovic, J., Knight, R.T., Endestad, T., and Helfrich, R. (2023). Subspace partitioning in the human prefrontal cortex resolves cognitive interference. Proc Natl Acad Sci U S A 120, e2220523120. 10.1073/pnas.2220523120.

54. Weber, J., Solbakk, A.-K., Blenkmann, A.O., Llorens, A., Funderud, I., Leske, S., Larsson, P.G., Ivanovic, J., Knight, R.T., Endestad, T., et al. (2024). Ramping dynamics and theta oscillations reflect dissociable signatures during rule-guided human behavior. Nat Commun 15, 637. 10.1038/s41467-023-44571-7.

55. Smith, S.M., and Nichols, T.E. (2009). Threshold-free cluster enhancement: addressing problems of smoothing, threshold dependence and localisation in cluster inference. Neuroimage 44, 83–98. 10.1016/j.neuroimage.2008.03.061.

56. Mensen, A., and Khatami, R. (2013). Advanced EEG analysis using threshold-free cluster-enhancement and non-parametric statistics. Neuroimage 67, 111–118. 10.1016/j.neuroimage.2012.10.027.

## References

1. Kerns, J. G. et al. Anterior Cingulate Conflict Monitoring and Adjustments in Control. Science 303, 1023–1026 (2004).

2. Kerns, J. G. Anterior cingulate and prefrontal cortex activity in an FMRI study of trial-to-trial adjustments on the Simon task. NeuroImage 33, 399–405 (2006).

3. Sheth, S. A. et al. Human Dorsal Anterior Cingulate Cortex Neurons Mediate Ongoing Behavioral Adaptation. Nature 488, 218–221 (2012).

4. Bartoli, E. et al. Temporal Dynamics of Human Frontal and Cingulate Neural Activity During Conflict and Cognitive Control. Cereb Cortex 28, 3842–3856 (2018).

5. Gratton, G., Coles, M. G. & Donchin, E. Optimizing the use of information: strategic control of activation of responses. J Exp Psychol Gen 121, 480–506 (1992).

6. Shenhav, A., Botvinick, M. M. & Cohen, J. D. The expected value of control: An integrative theory of anterior cingulate cortex function. Neuron 79, 217–240 (2013).

7. Chinn, L. K., Pauker, C. S. & Golob, E. J. Cognitive control and midline theta adjust across multiple timescales. Neuropsychologia 111, 216–228 (2018).

8. Lachaux, J.-P. et al. Estimating the time-course of coherence between single-trial brain signals: an introduction to wavelet coherence. Neurophysiol Clin 32, 157–174 (2002).

